# Molecular and physiological characterization of brassinosteroid receptor BRI1 mutants in *Sorghum bicolor*

**DOI:** 10.1101/2024.04.22.590590

**Authors:** Andrés Rico-Medina, David Blasco-Escámez, Juan B. Fontanet-Manzaneque, Natalie Laibach, Fidel Lozano-Elena, Damiano Martignago, Ana I. Caño-Delgado

## Abstract

**SUMMARY:**

- The high sequence and structural similarities between BRI1 brassinosteroid receptors of Arabidopsis (AtBRI1) and sorghum (SbBRI1) prompted us to study the functionally conserved roles of BRI in both organisms.
- Introducing sorghum SbBRI1 in Arabidopsis *bri1* mutants restores defective growth and developmental phenotypes to WT levels.
- Sorghum mutants for SbBRI1 receptors show defective BR sensitivity and impaired growth and development throughout the entire sorghum life cycle. Embryonic analysis of sorghum primary roots permit to trace back root growth and development to early stages, revealing the functionally conserved roles of SbBRI1 receptor in BR perception during meristem development. RNA-seq analysis uncovers the downstream regulation of the SbBRI1 pathway in cell wall biogenesis during cell growth.
- Together, these results uncover that sorghum SbBRI1 receptor protein play functionally conserved roles in plant growth and development, while encourage the study of BR pathways in sorghum and its implications for improving resilience in cereal crops.

## INTRODUCTION

Brassinosteroids (BRs) hormones are essential for plant growth and development and for the adaptation of plants to environmental stress (Planas-Riverola *et al*., 2019), endorsing their interest in crop improvement (Gupta *et al*., 2020). In *Arabidopsis thaliana* (Arabidopsis), BRs are perceived in the plasma membrane, via leucine-rich-repeat receptor-like-kinase (LRR-RLK) proteins of the BRI1 (BRASSINOSTEROID INSENSITIVE 1) family. Other BRI1-like homologues, namely BRI1-LIKE 1, 2 and 3 (BRL1/2/3) are present in Arabidopsis, from which only BRL1 and 3 can bind BR molecules with high affinity. Contrasting with the ubiquitous expression pattern in BRI1 (Li & Chory, 1997), BRL1 and BRL3 are enriched in the vasculature and stem cells (Caño-Delgado *et al*., 2004; Fàbregas *et al*., 2013; Salazar-Henao *et al*., 2016). BRs bind to BRI1-like receptors at a hydrophobic pocket known as “island domain”, created by a stretch of ∼70 amino acids that interrupt the LRR tandems (Kinoshita *et al*., 2005). The ligand binding creates a docking platform that favors BRI1 heterodimerization with the co-receptor BAK1 (Li *et al*., 2002; Hothorn *et al*., 2011; Bojar *et al*., 2014). This initiate a BR signaling cascade, that that triggers dephosphorylation and activation of transcription factors BRI1-EMS SUPRESSOR1 (BES1) and BRASSINAZOLE RESISTANT1 (BZR1) (Wang *et al*., 2002; Yin *et al*., 2002) by dephosphorylation and subsequent translocation of BES1 and BZR1 to the nucleus to control BR-regulated gene expression.

The last twenty years of molecular anaysis of Brassinosteroids in the roots have uncover that BR hormones operates with high spatio-temporal resolution (Planas-Riverola *et al*., 2019). Using the Arabidopsis root as a model, the important roles of BRs in the different cellular processes that govern root growth and development have been shown (González-García *et al*., 2011; Hacham *et al*., 2011). BRs promote root growth by controlling meristem development and the shift to cell elongation (Vilarrasa-Blasi *et al*., 2014; Pavelescu *et al*., 2018; Betegón-Putze *et al*., 2021; Nolan *et al*., 2023). Cell wall modifications during cell elongation are modulated by brassinosteroids in a cell-specific fashion (Li *et al*., 2021; Kelly-Bellow *et al*., 2023), yet the precise regulation of this process is not clear (Graeff *et al*., 2020). In the root meristem, BR hormones act in a paracrine way (Lozano-Elena *et al*., 2018) and can travel from cell to cell via plasmodesmata (PD), where they modulate its mobility by controlling its permeability (Wang *et al*., 2023).

Mutants defective in BR signaling or synthesis have been generated in other monocot crops, showing a characteristic stunted growth. A potent drive in the research for semi-dwarf mutants is their agronomical value as lodging resistant and more productive plants, as seen since the Green Revolution (Salas Fernandez *et al*., 2009). The semi-dwarf phenotype of the recessive *uzu* barley gene, used by Japanese breeders since the 1930s, is caused by a missense mutation in the kinase domain of HvBRI1 (Chono *et al*., 2003). Other barley dwarf mutants are allelic mutants of HvBRI1, or BR-deficient mutants with altered functions of three BR biosynthetic genes (Dockter *et al*., 2014). In maize, where BR-insensitive mutants have never been found in germplasm collections, compromising BRI1 homologs function with RNAi results in various degrees of dwarfism in the transgenic lines and diminished root growth inhibition by BL (Kir *et al*., 2015). The two classical dwarf maize *na1* and *brd1* carry mutations in BR biosynthetic genes (Hartwig *et al*., 2011; Makarevitch *et al*., 2012). In rice, most of the homologs of Arabidopsis BR biosynthetic genes have been characterized, with BR-deficient mutants also presenting reduced height and internode elongation (Castorina & Consonni, 2020). A similar phenotype was observed in the *d61* dwarf line, an OsBRI1 mutant that presents reduced root sensitivity to brassinolide (BL) treatments (Yamamuro *et al*., 2000).

*Sorghum bicolor* (sorghum) is a monocotyledon C4 cereal closely related to maize and sugarcane that ranks as the fifth most cultivated crop in the world (*FAOSTAT statistical database*., 2020; Chadalavada *et al*., 2021), emerging also as a bioenergy-producing crop in soils not fit to food-crop growth (Rocateli *et al*., 2012). Due to its resistance to drought and elevated temperatures, it is extensively cultivated in arid areas of the planet. The diploid sorghum genome presents minimal gene redundancy (Paterson *et al*., 2009), and its small size and complete sequence coverage allows for amenable functional genomics for other related crops such as maize or sugarcane (Swigoňová *et al*., 2004; Hughes *et al*., 2014). Still, it remains highly recalcitrant to genetic transformation reporting very low rates of success (O’Kennedy *et al*., 2006; Raghuwanshi & Birch, 2010). Despite sorghum potential for climate resilient agriculture, BR signaling components remain understudied, although recently there have been some efforts in this direction (Yamaguchi *et al*., 2016; Blasco-Escámez *et al*., 2017; Hirano *et al*., 2017).

Here, we report the functional characterization of brassinosteroid receptor kinase BRI1 in sorghum (SbBRI1). Driving the expression of SbBRI1 under the promoter 35S, the developmental defects caused by a *knockdown* of BRI1 (*bri1-301*) in Arabidopsis were restored to WT levels. Genetic and physiological analysis of two sorghum alleles carrying BRI1 mutations, SbBRI1-Pro407Leu (*ems87*) and SbBRI1-Val403Met (*ems72*) indicate conserved roles for SbBRI1 in promoting growth and development. Embryonic analysis of Sorghum root reveals the SbBRI1 receptor roles in meristem cell division and stem cell maintenance at the root apex. Collectively, our study unveil functionally conserved roles for SbBRI1 in promoting plant growth and development, while setting a solid foundation for the cellular analysis of roots in sorghum.

## RESULTS

### BRI1 receptor sequence conservation in *Sorghum bicolor*

To address the conservation of the BRI1 protein sequence among land plant species, a phylogenetic tree was built based on ortholog protein sequences (Fig. 1a). SbBRI1 is closely related to maize BRI1 (*Zea mays*) (93.6% of identity) and other monocot crops such as barley or wheat (*Hordeum vulgaris* or *Triticum aestivum)* (79.6% and 79.2% respectively). Similar conservation in the extracellular domain can be also observed in other plant species. (Fig. S1a). AtBRI1 and SbBRI1 share a protein sequence identity of 52.6%. Both leucine-rich-repeats in the extracellular domain, intramembrane and kinase intracellular domains are strongly conserved. The majority of differences are found at the extracellular domain, where SbBRI1 has only 19 LRRs at the N-terminal in comparison to the 24 LRRs found in AtBRI1 (Fig. 1b, S1b). The high degree of similarity in the kinase intracellular region suggests functional conservation during evolution (Ferreira-Guerra et al., 2020). In addition, structure modelling of the extracellular domain based on Arabidopsis BRI1 crystal (PDB: 3RJ0 (Hothorn *et al*., 2011)) reveals a high degree of structural similarity between both BRI1 proteins (Fig. 1b), keeping the horseshoe-like structure and the island domain highly conserved. The amino acidic substitutions of the two independent alleles carrying BRI1 mutations used in this study are shown in purple (Fig. 1b), which fall on a conserved LRR region in the extracellular domain of the protein. Furthermore, the conservation of the island domain, critical for hormone binding (Kinoshita *et al*., 2005), suggests a conservation of the protein function as BR receptor.

**Figure 1.**
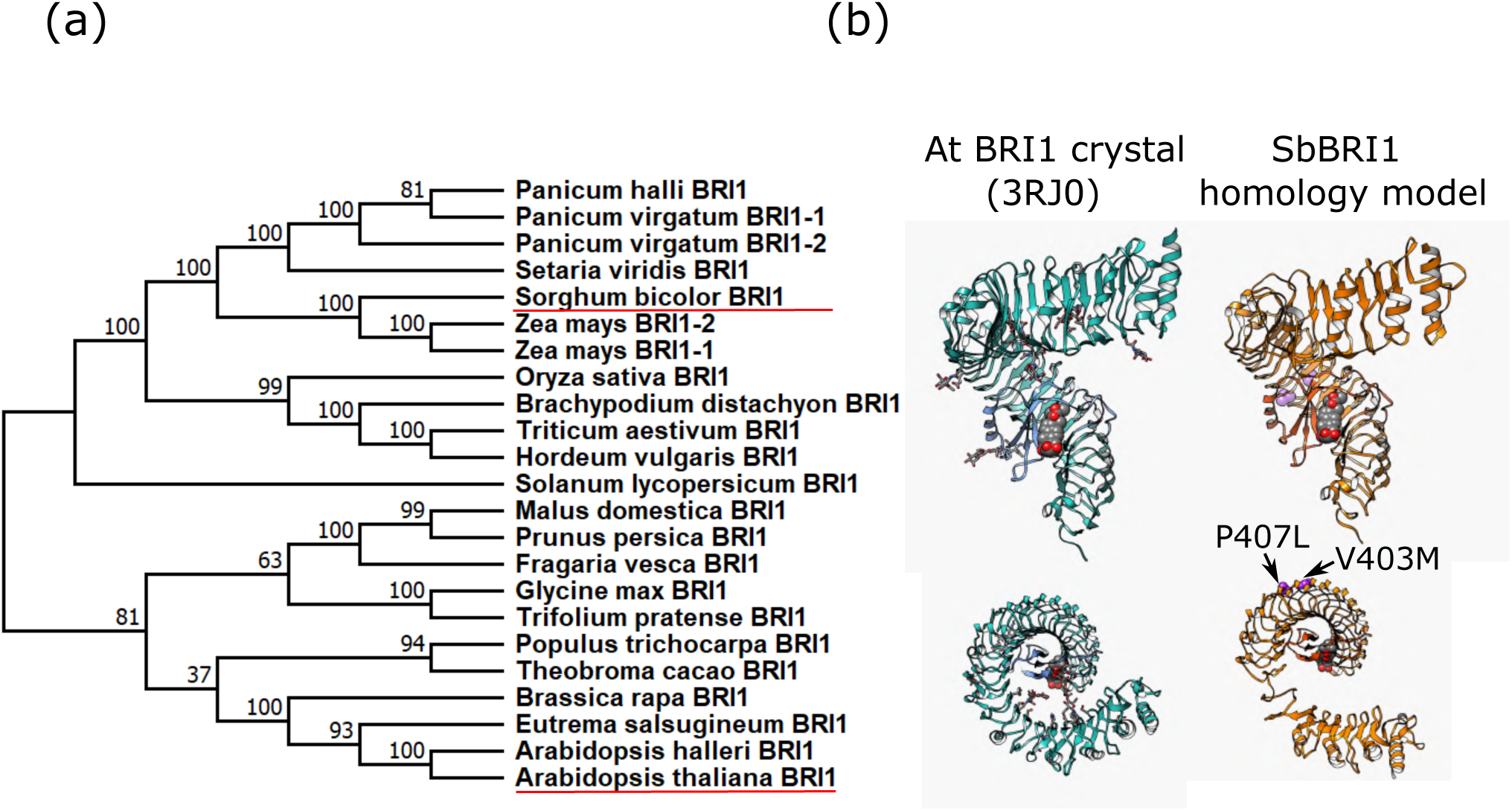
BRI1 structure is conserved in Sorghum bicolor. **a)** Sorghum BRI1 is more closely related to BRI1 from *Zea mays* and other crops than to Arabidopsis thaliana (both underlined in red). **b)** *In silico* 3D modelling of the LRR region of both Arabidopsis and SbBRI1 reveals a great conservation in structure, including the island domain, shown together with BL. Both studied EMS alleles are shown.

### Sorghum BRI1 protein restore *bri1* mutant phenotypes of Arabidopsis

To analyze the functional conservation of SbBRI1, a 35S:SbBRI1-GFP construct was introduced in Arabidopsis *bri1-301* mutants (35S:SbBRI1-GFP; *bri1-301*, hereinafter named SbBRI*ox; bri1-301*, see Methods). Adult homozygous SbBRI1*ox; bri1-301* plants exhibited a WT phenotype, in terms of overall size, rosette structure, and flowering time (Fig. 2a, Fig. S2a), restoring the dwarf phenotype of Arabidopsis *bri1-301* mutant (Kang *et al*., 2010), supporting that SbBRI1 can function as a BR receptor in promoting plant growth similarly to AtBRI1.

**Figure 2.**
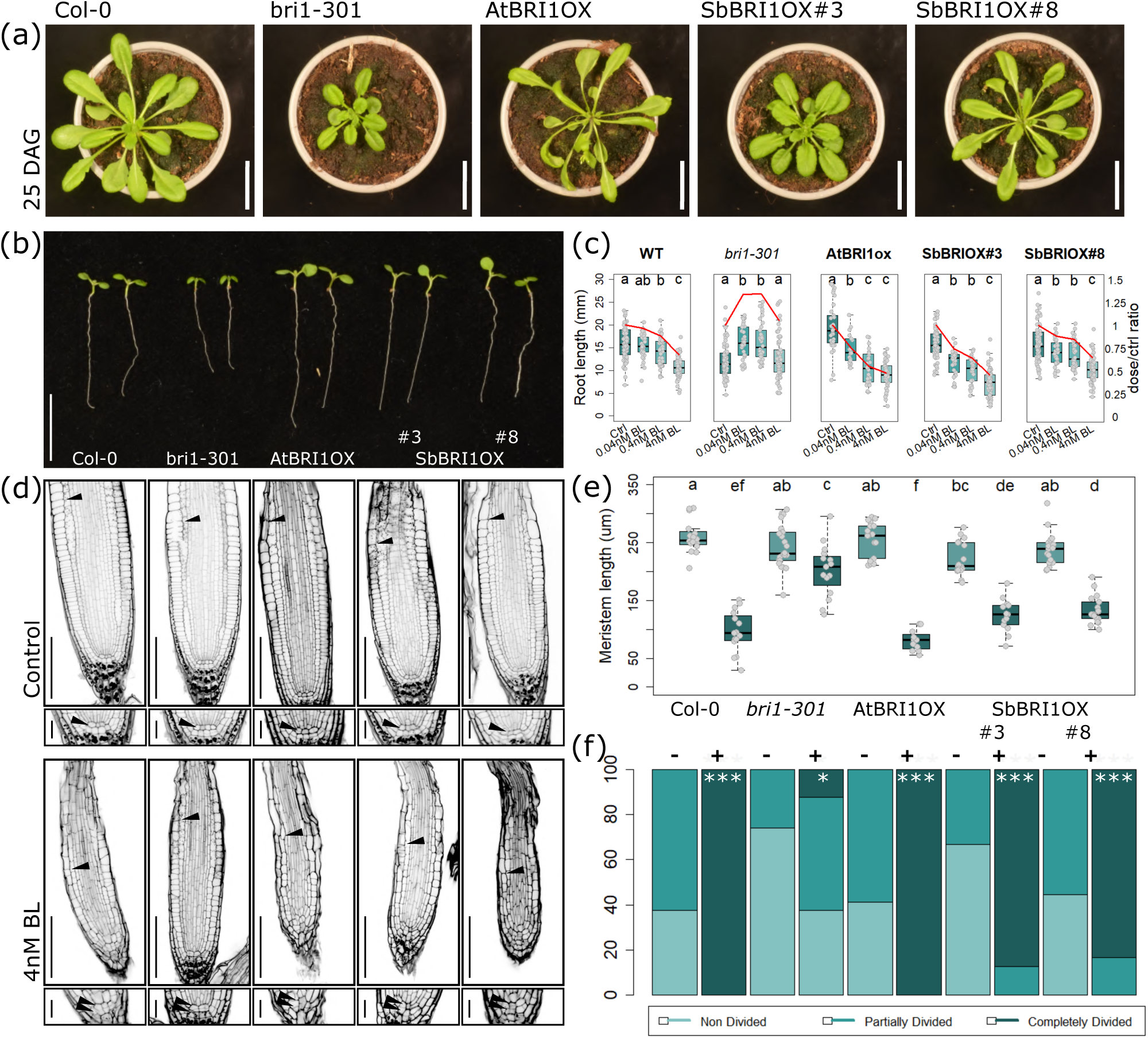
SbBRI1 can complement its AtBRI1 mutant counterpart phenotypes at macro and microscopical levels. **a)** 35S:SbBRI1-GFP; *bri1-301* (SbBRI1*ox*) can complement adult Arabidopsis *bri1-301* phenotype. White bars represent 4cm. **b, c)** SbBRI1*ox* complements root length phenotype of Arabidopsis *bri1-301* in control conditions. **(b)** and after BL supplementation **(c)**. White bar represents 1cm. **d, e, f)** SbBRI1ox complements meristem size **(e)** and QC division phenotype **(f)** of Arabidopsis *bri1-301* in control conditions and after BL supplementation. Black bars in **(d)** represents 100µm in root tip panels and 20µm in small QC panels.

The primary root of SbBRI1*ox*; *bri1-301* is longer than *bri1-301* roots, similar to WT (Fig. 2b). Upon BL application, a dose-dependent reduction in root length was observed in WT and AtBRI1 overexpressor plants (35S:BRI1-GFP; Col-0, herein named AtBRI*ox*), whereas *bri1-301* mutants were insensitive to BRs (Fig. 2c; González-García et al., 2011). The roots of SbBRI1*ox; bri1-301* plants behaved similarly to AtBRI1*ox* and WT (Col-0), of Arabidopsis (Fig. 2b), thus rescuing *bri1-301* phenotype of BL insensitivity. Upon BL application the root meristem length (Fig. 2d, e) and QC division rate (Fig. 2d, f) of SbBRI*ox; bri1-301* plants were restored to WT levels.

To further substantiate sorghum SbBRI1 functionality in Arabidopsis, we analyzed its ability to activate the downstream BR signaling pathway by assessing the phosphorylation status of BES1 in SbBRI1*ox; bri1-301* plants. Upon BL application, SbBRI1 was found to dephosphorylate BES1 protein in WT Col-0 and similarly in AtBRI1*ox; bri1-301* lines (Fig. S2b). Together, our results indicate that SbBRI1 can function as BR receptor activating downstream effectors in the pathway similar to AtBRI1.

### Characterization of *bri1* receptor mutants in sorghum

To investigate the native role of BRI1 receptor kinase in sorghum, mutant lines for SbBRI1 from an EMS-mutagenized population of BTx623 variety (Jiao *et al*., 2016) were identified. Two different SbBRI1 homozygous mutants were selected, *ems72* (V403M) and *ems87* (P407L) both possessing a point mutation in the extracellular LRR domain of the SbBRI1 receptor (Fig. 1b). To reduce the number of background mutations, two rounds of genetic backcrosses were performed using the non-mutagenized BTx623 variety as parental (Fig. S6, see Methods). Subsequent experiments were done by comparing the backcrossed *bri1* mutants for SbBRI1 with their SbBRI1 WT direct siblings, herein named *bri1-ems72* and WT-*ems72*.

In greenhouse conditions, mature *bri1-ems72* plants showed a reduction in overall plant growth, (Fig. 3a) during all developmental stages until maturity (Fig. 3b, c, d). Mature plants of *bri1*-*ems72* and *87* exhibited shorter panicles and a significantly reduced grain yield (Fig. 3e, h, Figure S3a, b), that might be attributed to a reduced grain number and not a reduced grain size or weight (Fig. S3c, d, e). Overall, our results indicate that sorghum SbBRI1 has a promoting role in plant growth by controling organ size.

**Figure 3.**
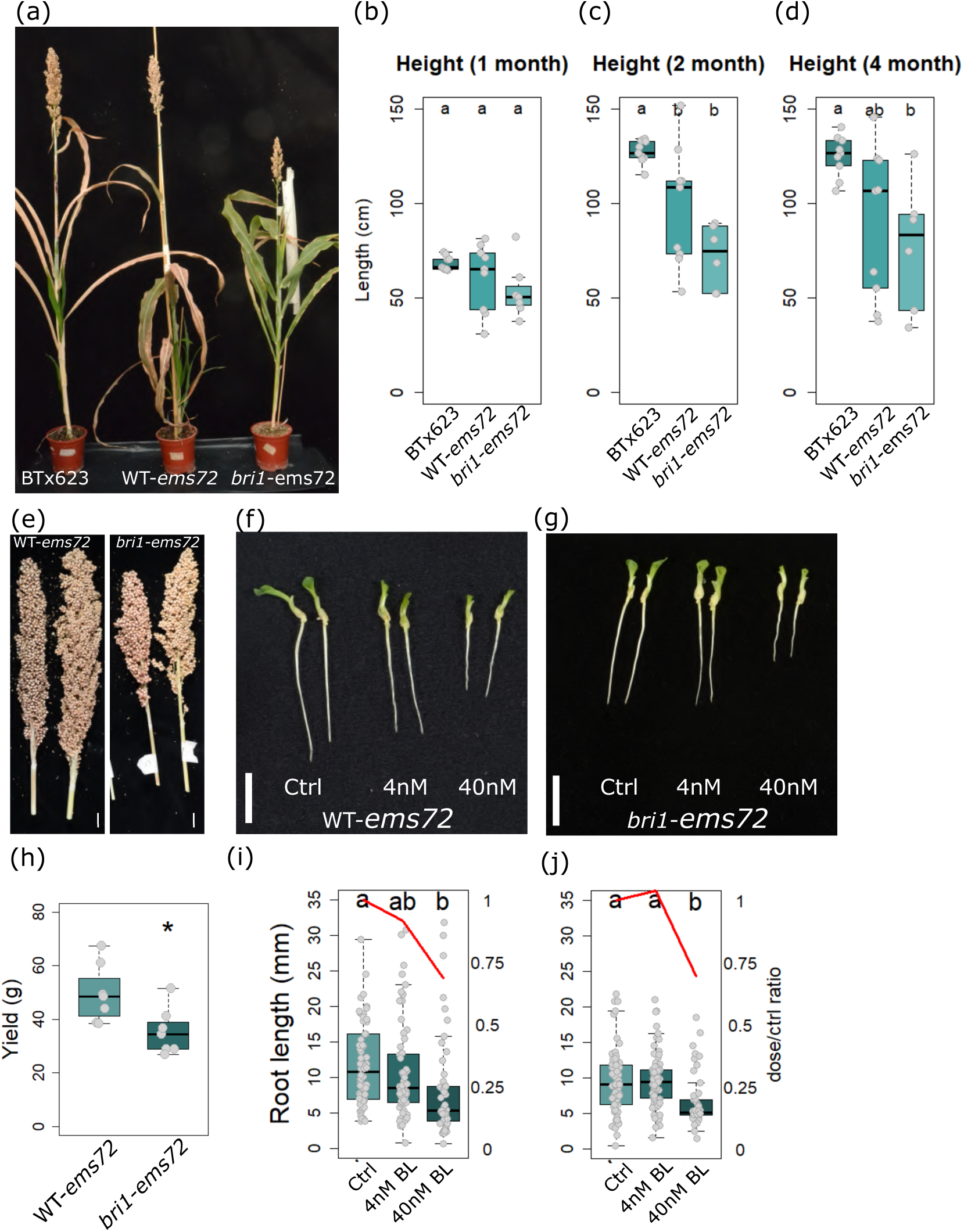
*bri1*-ems sorghum shows reduced plant height, yield, and reduced root length and a defective BL response. **a)** 4-month-old *bri1-ems72* plants display reduced plant height. **b, c, d)** *bri1-ems72* plants show reduced plant height during all developmental stages when compared against their WT counterparts and against WT control. **e)** *bri1-ems72* panicles present reduced yield when compared with their WT counterparts. Total yield weight is shown in **h). f, g, i, j)** Embryonic roots of *bri1*-*ems72* present reduced length and reduced sensitivity to BL supplementation when compared with their WT counterparts.

Next, explanted embryos grown *in vitro* were used to study the embryonic root development and standardize root growth analysis (see Methods). The embryonic roots of the WT sorghum BTx623 variety were found to germinate more homogeneously and be shorter than when using whole seeds (Fig. S4a), proving to be an amenable method of working in vitro with organs from non-model species and indicating that sorghum embryonic root can be used as a model in developmental studies. In control conditions, eight-day-old embryonic roots of *bri1-ems72* mutants were shorter than WT-*ems72* and showed reduced sensitivity to 4nM BL (Fig. 3g, h, k, l). Furthermore, when grown in a hydroponics setting, *bri1*-*ems87* also showed insensitivity to low BL concentrations (0.04nM and 0.4 nM of BL) (Fig. S4b, c). Together with the observed structural and functional conservation of SbBRI1 protein, genetic analysis shows the role of BRI1 receptor in sorghum BL perception and yield.

### Cellular anatomy of sorghum primary root in WT and SbBRI1 mutants

To further investigate the development of the primary root apex and the overall root architecture of *bri1* mutants in sorghum, mPS-PI staining protocol was adapted to embryonic roots from (Truernit *et al*., 2008; Kirschner *et al*., 2017) (Fig. 4). WT BTx623 roots shows a meristematic region of approximately fifteen cells, measured at the epidermis cell layer (Fig. 4a). In the medial plane, the stem cell niche was clearly visualized. The QC region and distally localized columella stem cells appear underneath, followed by the differentiated columella cells presenting their characteristically stained starch granules (Supplementary Video 1, Supplementary Video 2). Upon exogenous BL application, the meristem region size varies accordingly with described in Arabidopsis (González-García *et al*., 2011), and progressive disorganization was observed at the QC and stem cell region (Fig. 4b-e).

**Figure 4.**
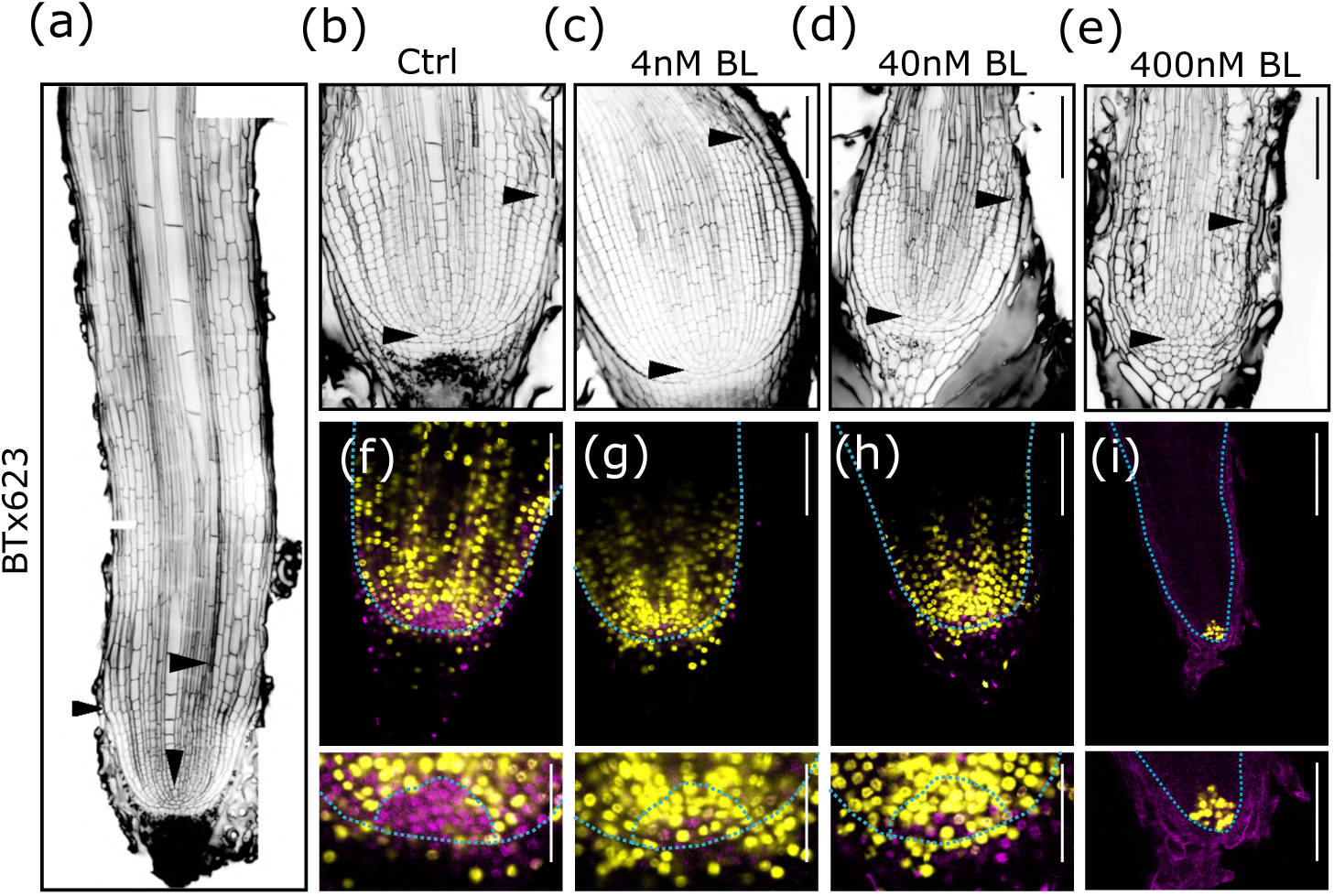
Root meristem architecture and BR response of BTx623 WT sorghum is shown by adapting common Arabidopsis confocal microscopy techniques. **a)** Root architecture of BTx623 after mPS-PI staining. From upper to lower, arrowpoints mark: xylem start, meristem region border, and QC region. **b, c, d, e)** Root meristem architecture of BTx623 grown at different BL concentrations. At increased BL amounts, Root thinning, meristem shrinkening and cell disorganization is observed. **f)** EdU staining of root meristem region of BTx623 sorghum reveals (in yellow) the cells dividing in a given timeframe (12h) and, by DAPI counterstaining, (in purple) an undivided set of cells that conforms the QC region. When grown at increased BL concentrations, **g, h, i)** EdU staining shows an increased cell division in the QC region **(g, h)** up to the point of total meristem exhaustion **(i)**. Black and white bars in **(b)** to **(i)** represent 80µm, while white bars in small QC panels represent 40µm.

To precisely define the QC region of the Sorghum root meristem, 5’-ethynyl-2’-deoxyuridine (EdU) cell proliferation assay was performed in embryonic roots (Fig. 4f-i). EdU is a thymidine analogue uptaken by the cell after growth media supplementation and incorporated in the genome of dividing cells, labelling it and thus allowing detection of said cells by confocal microscopy (in yellow). Together with all root nuclei counterstaining by DAPI (in purple), the technique allows the tracking of cell division in a given timeframe (12 hours, see Methods). In 8-day-old Sorghum WT BTx623 roots, the QC appears to be comprised of about 20 cells in its medial plane, and presents a semispherical shape under 3D imaging (Fig. 4f, Supplementary Video 1). Ever-increasing QC division was found under incremental BL supplementation, up to the point of root exhaustion at 400nM of BL, showing a significantly reduced cell division rate in an almost depleted meristem (Fig. 4i), accompanied by a severe cell disorganization of the stem cell region (Fig. 4e, i).

Upon BL application, WT-*ems72* shows rapid meristem shortening and cell disorganization (Fig. 5a, c), a response similar to that of BTx623, but more sensitive, probably due to background mutations. While *bri1*-*ems72* presents an overall similar response (Fig. 5b), meristem cell number at 4nM BL concentrations is significantly greater (Fig. 5c). No differences were found in meristem size response to BL (Fig. 5d). At 40nM BL concentration, there are not significant differences between genotypes and overall disorganization appears widely. These results also highlight the role of SbBRI1 in regulating root meristem cell division.

**Figure 5.**
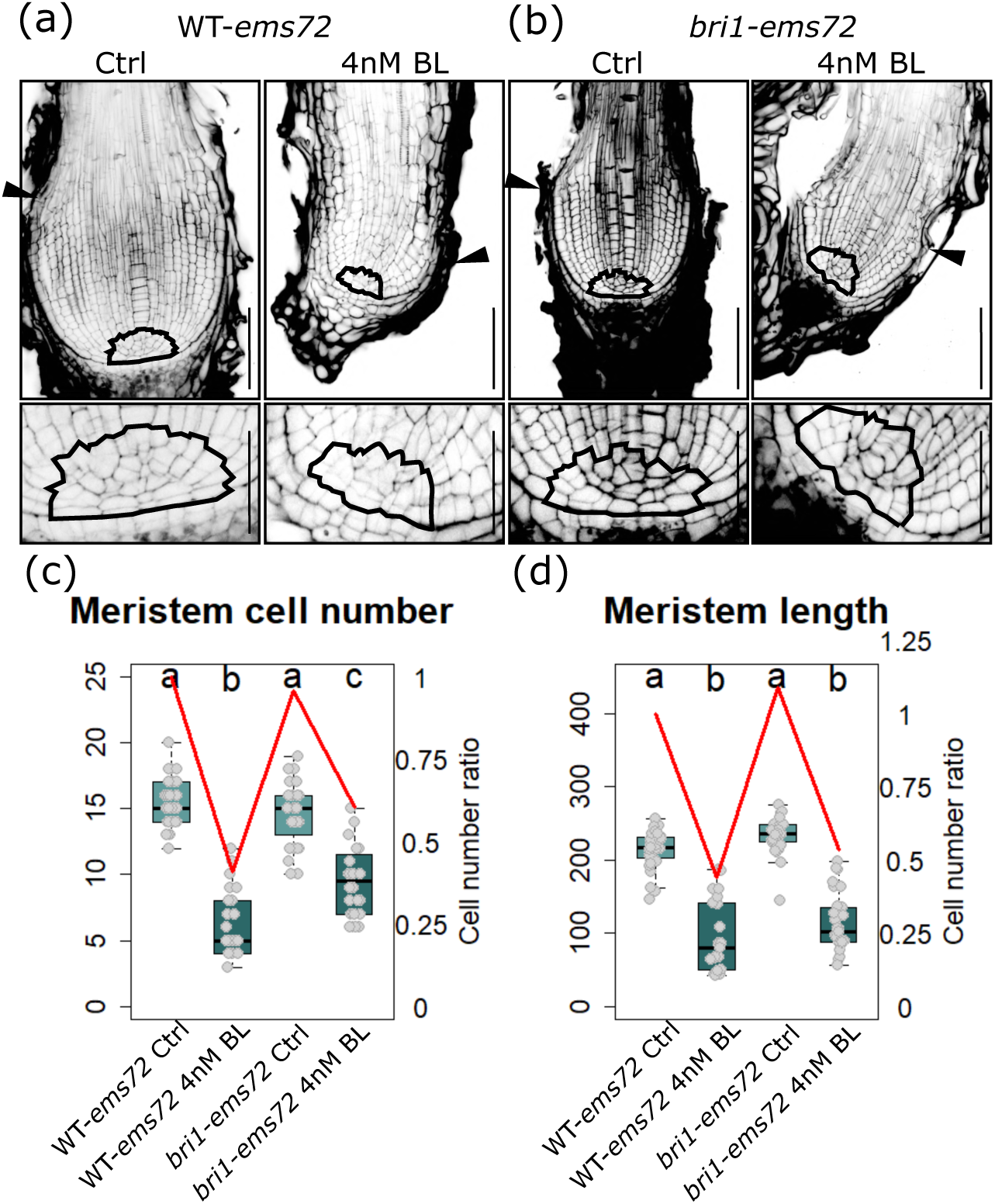
SbBRI1 is involved in root sensitivity of BRs. **a, b)** Root architecture of WT-ems72 and *bri1*-ems72 shows a reduced response of *bri1*-ems72 to BL supplementation. **c, d)** *bri1*-ems72 shows smaller cell number reduction than WT-ems72 when supplemented with BL **(c)** while the meristem length size reduction shows no significant differences **(d)**, fundamentally suggesting an impaired sensitivity to BL in *bri1*-ems72. Black bars in **(a)** and **(b)** root tip panels represent 80µm, while black bars in small QC panels represent 40µm.

In control conditions, *ems72* shows generally lesser cell division than BTx623 (Fig. 6). However, when supplemented with 4nM BL, overall cell division increases in the meristem (Fig. 6a, b, d, e), whereas *bri1*-*ems72* consistently shows a reduced cell division compared to its WT counterpart (Fig. 6g, h). At greater BL concentrations, the meristem region is usually exhausted, showing little to no meristematic cell division and a disorganized structure (Fig. 6c, f, Fig. S5a, b). Reduced BR sensitivity in *bri1*-*ems72* resulted in a reduction of the percentage of exhausted roots observed at 40nM BL (Fig. 6g, h). Our results uncover the roles of BRI1 receptor in Sorghum in meristem division and overall root growth, while opening embryonic roots analysis in sorghum at a cellular resolution to advance in our present understanding of root growth and development in cereal crops.

**Figure 6.**
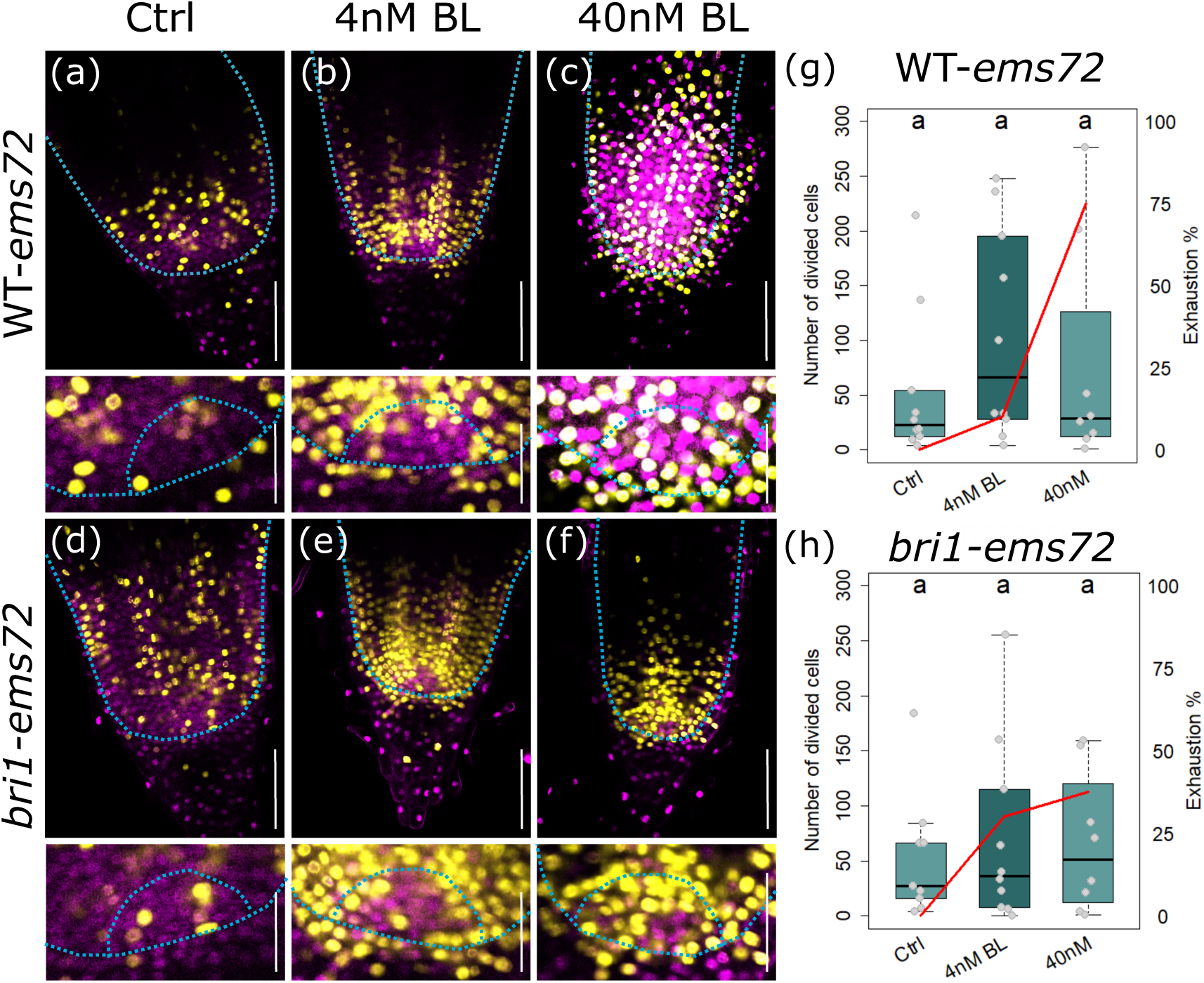
Confocal analysis of root meristem region revealed SbBRI1 role in meristem development. **a, b, c)** WT-*ems72* QC region presents increased cell division grown under increasing BL supplementation. **d, e, f)** *bri1*-*ems72* QC region presents increased cell division grown under increasing BL supplementation. **g, h)** A comparison of dividing cell number in WT-*ems72* and *bri1-ems72* reveals a reduced division rate in *bri1-ems72* at lower BL concentrations, together with a reduced *bri1-ems72* exhaustion rate at higher BL concentrations, fundamentally suggesting an insensitivity to BL due to the lack of a fully functional BRI1. White bars in root tip panels represent 80µm, while white bars in QC panels represent 40µm.

### Transcriptomic analysis shows conserved roles of SbBRI1 in cell wall loosening during cell elongation in sorghum

To study the role of BRI1 in sorghum in more molecular detail, RNA-seq analysis was used to investigate which processes are transcriptionally regulated by SbBRI1 in *bri1-ems72* roots against their sibling WT-*ems72*, (see Methods). Differential expression analysis revealed a limited number of deregulated genes (50 downregulated, 254 upregulated, Supplementary Table 4, Supplementary Table 5), probably because of transcriptional noise caused by the background mutations (FC>1.5, p value<0.05). GO enrichment analyses, based on 226 high confidence Arabidopsis orthologs of upregulated sorghum genes revealed enrichment in “plant-type cell wall loosening” (GO:0009828), “response to wounding” (GO:0009611), “cell wall macromolecule metabolic process” (GO:0044036), and “regulation of root development” (GO:2000280) categories (Fig. 7a), pointing towards the action of BRI1 on cell wall function. No enrichment nor BR-related transcripts (such as BRI1, BRL) were found deregulated among downregulated genes in our analysis. Gene deployment of the most significantly enriched GO categories (Fig. 7 b, c, d, e) show that the most deregulated ortholog genes per each category correspond to, respectively, expansin family members as AT1G20190 or AT1G65680 (EXPB2), chitinase enzymes as AT5G24090 (CHIA) or AT3G54420 (CHIV), fucosyltransferases as AT1G14100 (FUT8) and plasmodesmal-mediated primary root growth AT1G09560 (GLP5). As the sorghum genome is not as well annotated as Arabidopsis or other crop species, a search of the same deregulated sorghum genes reveals no different genes than using orthologs from Arabidopsis. Mostly expansins, chitinases and other enzymes are present. Together, our findings support the hypothesis that BRI1 functions of cell wall biosynthesis and remodeling are conserved in *Sorghum bicolor*.

**Figure 7.**
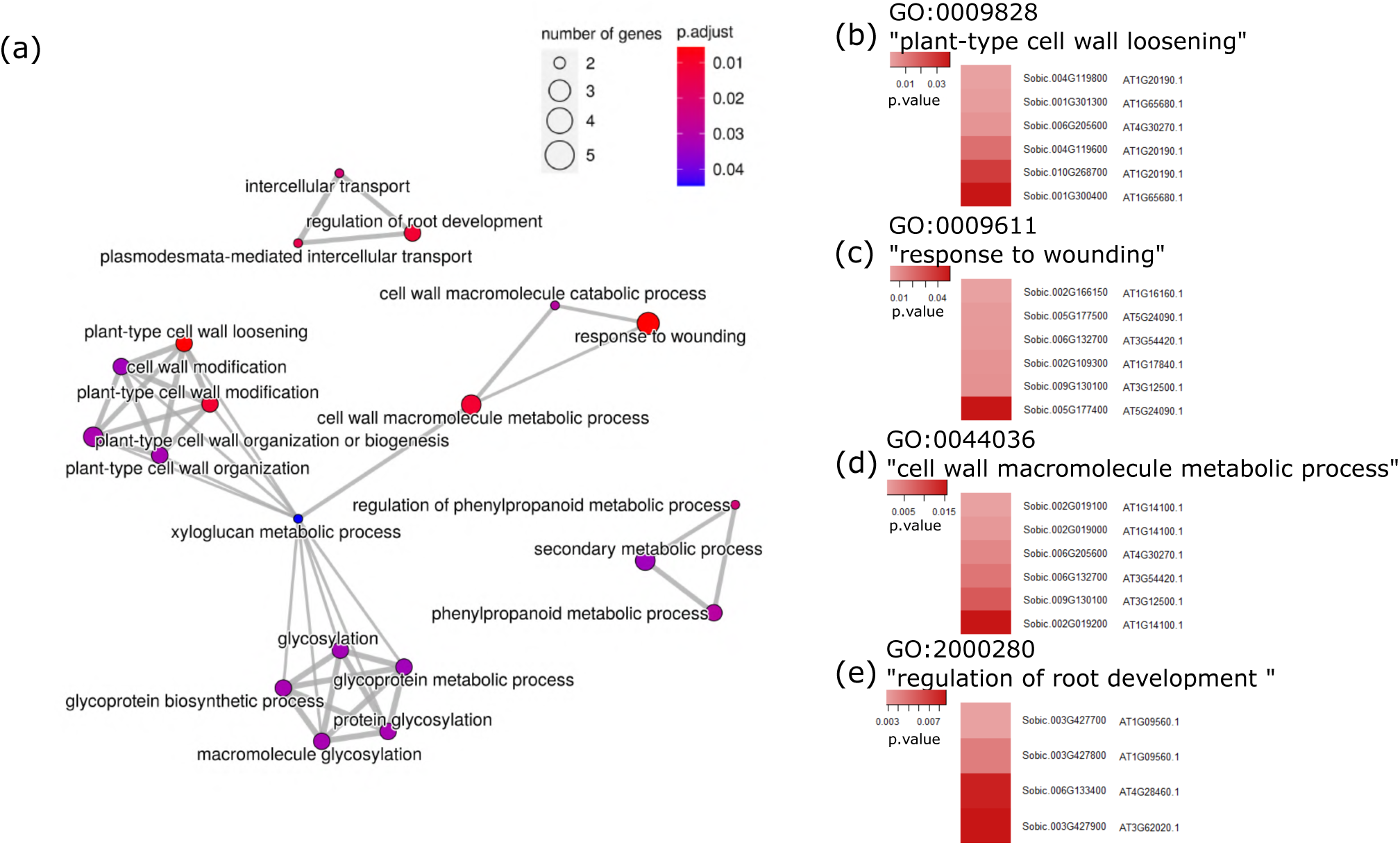
RNA-seq of *bri1vsWT* sorghum roots reveal BRI1 role in root growth. **a)** GOEA map of deregulated GO categories in sorghum *bri1-*ems72 *vs* WT-ems72 RNA-seq shows a role of SbBRI1 in cell wall metabolism regulation. **b), c), d) e)** Deployment of genes within the most deregulated GO terms from our upregulated set of genes. Sorghum gene name and closest Arabidopsis ortholog names are shown.

## DISCUSSION

In sorghum, there are two distinct LRR-RLs like AtBRI1, are the SbBRI1 and a putative SbBRL1, the later sharing highest homology to the BRI1-LIKE (BRL) family in Arabidopsis. Here, the genetic analysis of SbBRI1 mutants showing a reduced BR sensitivity conveys with similar previously described data for HvBRI1 and OsBRI1 mutants (Chono *et al*., 2003; Tong & Chu, 2012). The existing relationship between modified BRI1 receptor levels and proper seed and panicle development (Sun *et al*., 2021) is supportive of a direct relationship between BR biosynthesis and yield (Zhang *et al*., 2014; Singh *et al*., 2016). In Sorghum, SbBRI1 shares high amino acid sequence identity(52.6%) to AtBRI1 and similar three-dimensional structure including the island domain critical for BL binding (Kinoshita *et al*., 2005) and activation of the BR-mediated phosphorylation cascade (Z. Wang et al., 2002; Yin et al., 2002). Our data showing sorghum BRI overexpression in the Arabidopsis *bri1* receptor mutants resulting in phenotypes similar to WT to AtBRI1 overexpression (Wang *et al*., 2001) conveys on the functional conservation of the BR receptors in plants.

The meristem development phenotypes of SbBRI1 receptor mutants allied with the ones reported for *bri1* mutants of Arabidopsis (González-García *et al*., 2011; Lozano-Elena *et al*., 2018). The roles of Arabidopsis BRI1 in cell wall elongation (González-García *et al*., 2011; Li *et al*., 2021) and biosynthesis (Vert *et al*., 2005; Sun *et al*., 2010; Wolf *et al*., 2012) are similar to the ones identified in this study for SbBRI1, further suggesting the receptor function conservation during evolution (Ferreira-Guerra *et al*., 2020). Whether other key BR-signaling pathway components, such as BAK1 or BES1/BZR1, are at least partially conserved in sorghum is yet to be determined.

The BTx623 sorghum variety used in this study, that has been widely used in agriculture, presents a spontaneous mutation in the gene DW1, historically traced back to 1905 (R.E. Karper & Quinby, 1946). DW1 encodes a protein that positively regulates BR signaling by inhibiting BIN2 activity (Hirano *et al*., 2017) and causes partial dwarfism, and therefore increased lodging resistance and improved crop productivity (Hilley *et al*., 2016). When working with a gene theorized to be upstream of this described mutation, such as SbBRI1, it can be expected that the observed phenotypes be mitigated by other mutations such as the one present in BTx623. Nonetheless, our study shows that a compromised BRI1 function causes defective growth in all plant stages in sorghum, comparable with the ones reported in other species, clearing a path for future crop studies regarding brassinosteroids and developmental or stress-related processes (Vriet *et al*., 2012).

The analysis of roots in crop plants has been challenging due to size and/or cell wall composition, which increases the difficulty of a deep analysis like it is routinely performed in Arabidopsis roots. Obtaining good resolution imaging at a cellular level in cereals is challenging. Here, the use of freshly explanted sorghum embryos as a source material allowed to synchronize the germination, standardize *in vitro* growth and focus the study on the primary root, which arises from the scutellar node of the embryo (Hochholdinger *et al*., 2004). Embryonic root in maize supports plant growth by itself (Hetz *et al*., 1996), as we proved to be sufficient in sorghum. Our work provides adaptations for mPS-PI and EdU staining protocols for sorghum roots, allowing highly precise visualization at the cell-specific level, such as presented in (Kirschner *et al*., 2017) for barley. Routinely implementing these cell biology tools can be a turning point in our approach to cell-level studies of tissues in non-model plant organisms. In most crops, studies with root systems have earnt less attention as compared to the ones focused on aerial plant organs. Given the importance of the root for overall plant growth and adaptation to the changing environment (including BR pathway and receptors), the advent of studies highlighting the importance of plant hormones for root development and adaptation to abiotic stress resistance (Gupta *et al*., 2020) might help to generate greater-yielding cereals, by delving into the knowledge of tissue-specific plant responses to drought (Fàbregas *et al*., 2018; Gupta *et al*., 2023).

Together, these results highlight the importance of BR hormone signaling pathways in plant growth and development across phyla, and its implications in crop genetic improvement (Martignago *et al*., 2020). While the complexities of the molecular plant responses to drought are still being deciphered in Arabidopsis, transferring the current knowledge to crops of key importance for global food security will be central in years to come.

## MATERIALS AND METHODS

### Informatic tools and modeling

BRI1 alignment of different species was generated using Geneious Prime software v.2020.0.4. (https://geneious.com). SbBRI1 ectodomain structure was modelled based on the AtBRI1 ectodomain crystal structure bound to BL (PDB:3RJ0) using Modeller v9.23 (Sali, s. f.). Representation of the model and mapping of the mutated residues were generated with UCSF Chimera (Pettersen *et al*., 2004)

### Plant material

Arabidopsis seedlings, ecotype Columbia, were sterilized with 35% bleach in deionized, sterile water (dH2O), plated in MS0.5-media, and grown vertically at 22°C in long day conditions for 6 days.

For sorghum whole seedlings, seeds were sterilized with 35% bleach in deionized sterile water for 10 minutes, washed five times in dH2O, and embedded overnight at 4°C in dark. Then, seeds were placed in MS0.5- and grown at 28°C for 8 days before imaging.

BTx623 variety sorghum panicles between 13-17 days post anthesis were cut from the plant and sterilized with 50% bleach in deionized sterile water with 0.5% Triton X-100 for 10 minutes with agitation, and then washed 5 times with dH2O before embryo extraction. Embryos were then plated in MS0.5-media, and grown vertically for 8 days at 28°C, in 12/12 light conditions. For RNA-seq, sorghum BC2F2 embryos of ems72 line were grown as explained and individually collected at 8 days after germination. Aerial organs were also individually collected for PCR genotyping of *bri1* (Fig. S6). For each biological replicate, a minimum of six *bri1-ems72* and WT-*ems72* roots were pooled to homogenize as much as possible the genetic background of the samples. For sorghum phenotyping, the same strategy was followed, however, BC2F2 plants were genotyped in the same way and their homozygous BC2F3 offspring was used to perform the experiments. A graphical methodology can be found in Fig. S6, genotyping primers can be found in Supplementary Table 1.

### Complementation of *bri1* mutants in Arabidopsis

Sorghum BRI1 coding sequence was cloned via Gateway Recombination Cloning Technology (Invitrogen) to obtain pEN-L1L2-SbBRI1 entry clone. By LR recombination reaction with pGWB406 (Nakagawa *et al*., 2007) together with p4P1r-35S, the destination vector p35S:SbBRI1-GFP was obtained. Arabidopsis *bri1-301* plants were transformed by floral dip to introduce the p35S:SbBRI1-GFP vector, and stable lines carrying single copy of the transgene were selected for this study. Supplementary Table 3 includes used primer sequences and Fig. S7 shows the plasmid maps.

### Western Blotting

Total protein was extracted from 6 days old Arabidopsis seedlings and resolved by SDS-PAGE. Western Blot analysis was performed as described in (Fàbregas et al., 2013) using primary antibodies against BES1 (kindly provided by Prof. Yanhai Yin) H3, (Yin *et al*., 2002) and GFP (Sigma-Aldrich) and anti-rabbit secondary antibody IgG (GE Healthcare UK).

### Microscopy

6-day-old Arabidopsis seedlings and 8-day-old root tips cut from sorghum embryos were submit to mPS-PI protocol, following Truernit et al., 2008. Standard protocol, as described, was used for Arabidopsis seedlings. For sorghum roots, protocol for floral stalks was followed with the following modifications: 50% methanol, 10% acetic acid fixation time was extended to 48 hours minimum, 1% periodic acid fixation was carried out under vacuum conditions for a minimum of 1 hour, and Schiff reagent with propidium iodide incubation was carried out under agitation for at least 3 hours to ensure staining of the more internal cell layers of the root tip. After chloral hydrate clearing, stained roots were mounted in excavated slides using Hoyer’s media, left to dry and observed using an Olympus FV1000 confocal microscope (20X and 40X objective, oil immersion, laser excitation set to 536-617nM). Images were taken in the medial plane of the root and different data was measured with ImageJ software.

For EdU staining, 7 and a half-day-old sorghum seedlings were transferred from 0.5MS-with vitamins media to plates of the same media, also containing 10µM EdU in DMSO (plus 400nM BL for the treated samples) or the corresponding amount of DMSO as mock and grown in that media for 12 hours. For the staining protocol we followed (Kirschner *et al*., 2017) staining of barley roots with a few modifications: root tips were cut and fixated for at least one hour under vacuum, permeabilized for another hour under vacuum, and submitted to Click-it reaction under vacuum as per Click-It Edu Alexa Fluor 488 Imaging Kit (Thermofisher) including nuclei counterstaining. Samples were then submitted to mild fixation with paraformaldehyde and cleared for around 15-20 days following (Warner *et al*., 2014). Microscopy was performed using Olympus FV1000 confocal microscope (40X objective, oil immersion, laser excitation set to 495/519nM). Images were taken in the medial plane of the root and different data measured with ImageJ software. For Fig. 6a, laser intensity was modified for better visualization as this image is a collage of many confocal pictures of the same root.

### Measuring and statistics

For in vitro plate images, root length was measured using ImageJ software (https://imagej.nih.gov/ij/). For cell division measures in Figure 6, ITCN plugin for ImageJ was used in the image channel corresponding to EdU staining using whole image keeping constant internal parameters. All graphics were generated using RStudio default plotting tools. Statistics were calculated in RStudio using Agricolae package for Tukey’s HSD test, and online with a 2×3 Fisher’s test (Freeman & Halton, 1946; Soper, 2022)A p value of at least p<0.05 is used as a threshold in all statistics unless otherwise stated. All experiments were carried out using at least three independent biological replicates.

### Transcriptomic profiling analysis

For sorghum RNA-seq, RNA was extracted from sorghum roots using Plant Easy Mini Kit (Qiagen). Stranded libraries were prepared using TrueSeq Stranded mRNA KIT (Ilumina). Paired-end sequencing (2×100bp) was performed using NovaSeq6000 (Ilumina) at minimum depth of 45M. Raw reads were quality-checked using FastQC tool (v0.11.2), trimmed 13bp at the 3’ end quality-trimmed and filtered at a minimum quality score of 28 (Phred+33, minimum read length of 75bp) using trimmomatic software (v0.38). Filtered reads were mapped to *Sorghum bicolor* v3.0.1 genome release using HISAT2 (2.1.0) aligner and quantified at a gene level using FeatureCount software (v1.6.2) on *Sorghum bicolor* v3.1.1 annotation file (Retrieved form Phytozome). For differential expression analysis, low-abundant raw counts were filtered out and remaining counts normalized by library size and TMM. Pairwise differential expression analysis was performed using edgeR package (v3.34.1) in R (v4.1.1). Gene Ontology enrichment analysis and representations were performed in R using clusterProfiler package (v4.0.5) based on best Arabidopsis orthologs hits for differentially regulated sorghum genes and the org.At.tair.db annotation package (v3.13.0, TAIR annotation release from April 2021). Both, raw reads files and processed counts have been deposited at Gene Omnibus Expression (GEO) database with identifier GSEXXXXX. Enriched GO categories were obtained using Arabidopsis orthologs and double checked in PantherDB (Mi et al., 2019) and Morokoshi Transcriptomic Database (Makita *et al*., 2015)

## ACKNOWLEDGEMENTS

The authors thanks, Prof. Yanhai Yin (Iowa State Univ.) for kindly providing us with the anti-BES1 antibodies, and Dr. Mar Marqués-Bueno for technical and hands-on advice on Western Blot.

## AUTHOR CONTRIBUTION

AR-M, DM and AIC-D outlined the manuscript, AR-M and AIC-D wrote the manuscript. AR-M performed the microscopy analysis and most of the data analysis. AR-M and DM performed WB experiments and the complementation analysis. DM cloned SbBRI1 CDS. DB-E, NL and JBF-M performed the sorghum EMS mutant research in adult plants. FL-E analyzed the raw transcriptomic data. AIC-D designed the research. All authors reviewed and edited the manuscript.

## SUPPLEMENTARY INFORMATION

**Supplementary Figure 1: A)** Sorghum and Arabidopsis BRI1 share a common sequence of kinase, transmembrane and island domain, suggesting functional conservation. Extracelular domain is mostly conserved with the exception of five LRR domains. **B)** Alignment of multiple BRI1 sequences from different species showing a general conservation of the intracellular and extracellular domains.

**Supplementary Figure 2: A)** SbBRI1 complements rosette size and flowering time in *bri1-301* Arabidopsis. **B)** BES1 dephosphorilation after BL treatment in SbBRI1 complemented lines of Arabidopsis *bri1-301* shows a functional conservation of BRI1 in sorghum.

**Supplementary Figure 3: A)** *bri1-ems87* panicles present reduced yield when compared with their WT counterpart. **C)** Seed weight of 50 seeds of WT-*ems* and *bri1-ems* mutants. **D, E)** size comparison of seeds of WT-*ems* and *bri1-ems* mutants.

**Supplementary Figure 4: A)** Comparison of in vitro grown BTx623 from whole seedling vs from explanted embryo. **B)** Hydroponic grown sorghum seedlings show a insensitivity to BL of *bri1*-*ems87*. **C)** *bri1*-*ems87* sorghum root shows insensitivity to low concentrations of BL in terms of root length.

**Supplementary Figure 5: A)** Example at 40nm BL of root meristem exhaustion in WT-*ems72* and *bri1*-*ems72* (in yellow, dividing cells), **B)** and with bright field bottom channel. **C)** Example of exhausted WT-*ems72* and *bri1*-*ems72* roots using mPS-PI show a great cell architecture disorganization in the root tip. **D)** At elevated BL concentrations, no differences were found between WT-*ems72* and *bri1*-*ems72*.

**Supplementary Figure 6: Diagram of backcrossing process to select our working seed lines.** EMS-mutated M4 seeds of both *bri1* alleles from Jiao et al, 2016, were backcrossed against their unmutagenized parental BTx623 and PCR-genotyped for *bri1* mutations. Homozygous *bri1* line was selected for a second round of backcrossing and their progeny PCR-genotyped. F2 of this second backcross was PCR-genotyped using the aerial organs and used at seedling stage to perform RNA-seq of the root organ of both *bri1* and WT alleles of the BRI1 gene. Sibling progeny of both were grown and used to obtain F3 seeds for a comparative phenotypical analysis of BRI1.

**Supplementary Figure 7: A)** After BP reaction, SbBRI1 coding sequence was cloned into pEN-L1L2 plasmid, giving pEN-L1L2-SbBRI1. **B)** pEN-L1L2-SbBRI1 and the p4P1r-35S vector were LR-recombined with destination vector pGWB406 to obtain p35S:SbBRI1-GFP

## SUPPLEMENTARY FILES

**Supplementary Table 1. Primer sequences used for sequencing of bri1 alleles in Arabidopsis and sorghum.**

**Supplementary Table 2. Protein sequences used for alignments.**

**Supplementary Table 3: Primer sequences used for cloning.**

**Supplementary Table 4: List of upregulated genes in *bri1* vs WT RNA-seq.**

**Supplementary Table 5: List of downregulated genes in *bri1* vs WT RNA-seq.**

**Supplementary Video 1:** mPS-PI of sorghum root meristem reveals the root architecture of the stem region. At 60X, the QC region can be extrapolated by comparing with EdU staining.

**Supplementary Video 2:** mPS-PI of sorghum root meristem reveals the root architecture of the stem region at 40X.

**Supplementary Video 3:** EdU staining of sorghum root meristem shows an undividing region (QC) surrounded by stem cells. Z-stack obtained from 40X confocal microscopy.

## REFERENCES

Betegón-Putze I, Mercadal J, Bosch N, Planas-Riverola A, Marquès-Bueno M, Vilarrasa-Blasi J, Frigola D, Burkart RC, Martínez C, Conesa A, et al. 2021. Precise transcriptional control of cellular quiescence by BRAVO/WOX5 complex in *Arabidopsis* roots. Molecular Systems Biology 17.

Blasco-Escámez D, Lozano-Elena F, Fàbregas N, Caño-Delgado AI. 2017. The Primary Root of Sorghum bicolor (L. Moench) as a Model System to Study Brassinosteroid Signaling in Crops. In: Russinova E, Caño-Delgado AI, eds. Methods in Molecular Biology. Brassinosteroids. New York, NY: Springer New York, 181–192.

Bojar D, Martinez J, Santiago J, Rybin V, Bayliss R, Hothorn M. 2014. Crystal structures of the phosphorylated BRI 1 kinase domain and implications for brassinosteroid signal initiation. The Plant Journal 78: 31–43.

Caño-Delgado A, Yin Y, Yu C, Vafeados D, Mora-García S, Cheng J-C, Nam KH, Li J, Chory J. 2004. BRL1 and BRL3 are novel brassinosteroid receptors that function in vascular differentiation in *Arabidopsis*. Development 131: 5341–5351.

Castorina G, Consonni G. 2020. The Role of Brassinosteroids in Controlling Plant Height in Poaceae: A Genetic Perspective. International Journal of Molecular Sciences 21: 1191.

Chadalavada K, Kumari BDR, Kumar TS. 2021. Sorghum mitigates climate variability and change on crop yield and quality. Planta 253: 113.

Chono M, Honda I, Zeniya H, Yoneyama K, Saisho D, Takeda K, Takatsuto S, Hoshino T, Watanabe Y. 2003. A Semidwarf Phenotype of Barley uzu Results from a Nucleotide Substitution in the Gene Encoding a Putative Brassinosteroid Receptor. Plant Physiology 133: 1209–1219.

Dockter C, Gruszka D, Braumann I, Druka A, Druka I, Franckowiak J, Gough SP, Janeczko A, Kurowska M, Lundqvist J, et al. 2014. Induced Variations in Brassinosteroid Genes Define Barley Height and Sturdiness, and Expand the Green Revolution Genetic Toolkit. Plant Physiology 166: 1912–1927.

Fàbregas N, Li N, Boeren S, Nash TE, Goshe MB, Clouse SD, de Vries S, Caño-Delgado AI. 2013. The BRASSINOSTEROID INSENSITIVE1–LIKE3 Signalosome Complex Regulates *Arabidopsis* Root Development. The Plant Cell 25: 3377–3388.

Fàbregas N, Lozano-Elena F, Blasco-Escámez D, Tohge T, Martínez-Andújar C, Albacete A, Osorio S, Bustamante M, Riechmann JL, Nomura T, et al. 2018. Overexpression of the vascular brassinosteroid receptor BRL3 confers drought resistance without penalizing plant growth. Nature Communications 9: 4680.

*FAOSTAT statistical database*. 2020. Rome: Food and Agriculture Organization of the United Nations.

Ferreira-Guerra M, Marquès-Bueno M, Mora-García S, Caño-Delgado AI. 2020. Delving into the evolutionary origin of steroid sensing in plants. Current Opinion in Plant Biology 57: 87–95.

Freeman G, Halton J. 1946. Note on exact treatment of contingency, goodness of fit and other problems of significance. Biometrika 38: 141–149.

González-García M-P, Vilarrasa-Blasi J, Zhiponova M, Divol F, Mora-García S, Russinova E, Caño-Delgado AI. 2011. Brassinosteroids control meristem size by promoting cell cycle progression in *Arabidopsis* roots. Development 138: 849–859.

Graeff M, Rana S, Marhava P, Moret B, Hardtke CS. 2020. Local and Systemic Effects of Brassinosteroid Perception in Developing Phloem. Current Biology 30: 1626–1638.e3.

Gupta A, Rico-Medina A, Caño-Delgado AI. 2020. The physiology of plant responses to drought. Science 368: 266–269.

Gupta A, Rico-Medina A, Lozano-Elena F, Marqués-Bueno M, Fontanet JB, Fàbregas N, Alseekh S, Fernie AR, Caño-Delgado AI. 2023. Brassinosteroid receptor BRL3 triggers systemic plant adaptation to elevated temperature from the phloem cells. Plant Biology.

Hacham Y, Holland N, Butterfield C, Ubeda-Tomas S, Bennett MJ, Chory J, Savaldi-Goldstein S. 2011. Brassinosteroid perception in the epidermis controls root meristem size. Development 138: 839–848.

Hartwig T, Chuck GS, Fujioka S, Klempien A, Weizbauer R, Potluri DPV, Choe S, Johal GS, Schulz B. 2011. Brassinosteroid control of sex determination in maize. Proceedings of the National Academy of Sciences of the United States of America 108: 19814–19819.

Hetz W, Hochholdinger F, Schwall M, Günter F. 1996. Isolation and characterization of rtcs, a maize mutant deficient in the formation of nodal roots. The Plant Journal: 845– 857.

Hilley J, Truong S, Olson S, Morishige D, Mullet J. 2016. Identification of Dw1, a Regulator of Sorghum Stem Internode Length (PK Subudhi, Ed.). PLOS ONE 11: e0151271.

Hirano K, Kawamura M, Araki-Nakamura S, Fujimoto H, Ohmae-Shinohara K, Yamaguchi M, Fujii A, Sasaki H, Kasuga S, Sazuka T. 2017. Sorghum DW1 positively regulates brassinosteroid signaling by inhibiting the nuclear localization of BRASSINOSTEROID INSENSITIVE 2. Scientific Reports 7: 126.

Hochholdinger F, Park WJ, Sauer M, Woll K. 2004. From weeds to crops: genetic analysis of root development in cereals. Trends in Plant Science 9: 42–48.

Hothorn M, Belkhadir Y, Dreux M, Dabi T, Noel JosephP, Wilson IA, Chory J. 2011. Structural basis of steroid hormone perception by the receptor kinase BRI1. Nature 474: 467–471.

Hughes TE, Langdale JA, Kelly S. 2014. The impact of widespread regulatory neofunctionalization on homeolog gene evolution following whole-genome duplication in maize. Genome Research 24: 1348–1355.

Jiao Y, Burke JJ, Chopra R, Burow G, Chen J, Wang B, Hayes C, Emendack Y, Ware D, Xin Z. 2016. A Sorghum Mutant Resource as an Efficient Platform for Gene Discovery in Grasses. The Plant Cell: tpc.00373.2016.

Kang B, Wang H, Nam KH, Li J, Li J. 2010. Activation-Tagged Suppressors of a Weak Brassinosteroid Receptor Mutant. Molecular Plant 3: 260–268.

Kelly-Bellow R, Lee K, Kennaway R, Barclay JE, Whibley A, Bushell C, Spooner J, Yu M, Brett P, Kular B, et al. 2023. Brassinosteroid coordinates cell layer interactions in plants via cell wall and tissue mechanics. Science 380: 1275–1281.

Kinoshita T, Caño-Delgado A, Seto H, Hiranuma S, Fujioka S, Yoshida S, Chory J. 2005. Binding of brassinosteroids to the extracellular domain of plant receptor kinase BRI1. Nature 433: 167–171.

Kir G, Ye H, Nelissen H, Neelakandan AK, Kusnandar AS, Luo A, Inzé D, Sylvester AW, Yin Y, Becraft PW. 2015. RNA Interference Knockdown of BRASSINOSTEROID INSENSITIVE1 in Maize Reveals Novel Functions for Brassinosteroid Signaling in Controlling Plant Architecture. Plant Physiology 169: 826–839.

Kirschner GK, Stahl Y, Von Korff M, Simon R. 2017. Unique and Conserved Features of the Barley Root Meristem. Frontiers in Plant Science 8: 1240.

Li J, Chory J. 1997. A Putative Leucine-Rich Repeat Receptor Kinase Involved in Brassinosteroid Signal Transduction. Cell 90: 929–938.

Li Z, Sela A, Fridman Y, Höfte H, Savaldi-Goldstein S, Wolf S. 2021. Optimal BR signalling is required for adequate cell wall orientation in the Arabidopsis root meristem. Development.

Li J, Wen J, Lease KA, Doke JT, Tax FE, Walker JC. 2002. BAK1, an Arabidopsis LRR Receptor-like Protein Kinase, Interacts with BRI1 and Modulates Brassinosteroid Signaling. Cell 110: 213–222.

Lozano-Elena F, Planas-Riverola A, Vilarrasa-Blasi J, Schwab R, Caño-Delgado AI. 2018. Paracrine brassinosteroid signaling at the stem cell niche controls cellular regeneration. Journal of Cell Science: jcs.204065.

Makarevitch I, Thompson A, Muehlbauer GJ, Springer NM. 2012. Brd1 Gene in Maize Encodes a Brassinosteroid C-6 Oxidase (S-B Wu, Ed.). PLoS ONE 7: e30798.

Makita Y, Shimada S, Kawashima M, Kondou-Kuriyama T, Toyoda T, Matsui M. 2015. MOROKOSHI: Transcriptome Database in Sorghum bicolor. Plant and Cell Physiology 56: e6–e6.

Martignago D, Rico-Medina A, Blasco-Escámez D, Fontanet-Manzaneque JB, Caño-Delgado AI. 2020. Drought Resistance by Engineering Plant Tissue-Specific Responses. Frontiers in Plant Science 10: 1676.

Nakagawa T, Suzuki T, Murata S, Nakamura S, Hino T, Maeo K, Tabata R, Kawai T, Tanaka K, Niwa Y, et al. 2007. Improved Gateway Binary Vectors: High-Performance Vectors for Creation of Fusion Constructs in Transgenic Analysis of Plants. Bioscience, Biotechnology, and Biochemistry 71: 2095–2100.

Nolan TM, Vukašinović N, Hsu C-W, Zhang J, Vanhoutte I, Shahan R, Taylor IW, Greenstreet L, Heitz M, Afanassiev A, et al. 2023. Brassinosteroid gene regulatory networks at cellular resolution in the *Arabidopsis* root. Science 379: eadf4721.

O’Kennedy MM, Grootboom A, Shewry PR. 2006. Harnessing sorghum and millet biotechnology for food and health. Journal of Cereal Science 44: 224–235.

Paterson AH, Bowers JE, Bruggmann R, Dubchak I, Grimwood J, Gundlach H, Haberer G, Hellsten U, Mitros T, Poliakov A, et al. 2009. The Sorghum bicolor genome and the diversification of grasses. Nature 457: 551–556.

Pavelescu I, Vilarrasa-Blasi J, Planas-Riverola A, González-García M, Caño-Delgado AI, Ibañes M. 2018. A Sizer model for cell differentiation in *Arabidopsis thaliana* root growth. Molecular Systems Biology 14.

Pettersen E, Goddard T, Huang C, Couch G, Greenblatt D, Meng C, Ferrin T. 2004. USCF Chimera - a visualization system for exploratory research and analysis.

Planas-Riverola A, Gupta A, Betegón-Putze I, Bosch N, Ibañes M, Caño-Delgado AI. 2019. Brassinosteroid signaling in plant development and adaptation to stress. Development 146: dev151894.

Raghuwanshi A, Birch RG. 2010. Genetic transformation of sweet sorghum. Plant Cell Reports 29: 997–1005.

R.E. Karper, Quinby JR. 1946. The history and evolution of Milo in the United States. Journal of the american society of agronomy.

Rocateli AC, Raper RL, Balkcom KS, Arriaga FJ, Bransby DI. 2012. Biomass sorghum production and components under different irrigation/tillage systems for the southeastern U.S. Industrial Crops and Products 36: 589–598.

Salas Fernandez MG, Becraft PW, Yin Y, Lübberstedt T. 2009. From dwarves to giants? Plant height manipulation for biomass yield. Trends in Plant Science 14: 454–461.

Salazar-Henao JE, Lehner R, Betegón-Putze I, Vilarrasa-Blasi J, Caño-Delgado AI. 2016. BES1 regulates the localization of the brassinosteroid receptor BRL3 within the provascular tissue of the Arabidopsis primary root. Journal of Experimental Botany 67: 4951–4961.

Singh A, Breja P, Khurana JP, Khurana P. 2016. Wheat Brassinosteroid-Insensitive1 (TaBRI1) Interacts with Members of TaSERK Gene Family and Cause Early Flowering and Seed Yield Enhancement in Arabidopsis (PK Trivedi, Ed.). PLOS ONE 11: e0153273.

Soper D. 2022. Fisher’s exact test calculator for a 2×3 contingency table.

Sun Y, Fan X-Y, Cao D-M, Tang W, He K, Zhu J-Y, He J-X, Bai M-Y, Zhu S, Oh E, et al. 2010. Integration of Brassinosteroid Signal Transduction with the Transcription Network for Plant Growth Regulation in Arabidopsis. Developmental Cell 19: 765–777.

Sun H, Xu H, Li B, Shang Y, Wei M, Zhang S, Zhao C, Qin R, Cui F, Wu Y. 2021. The brassinosteroid biosynthesis gene, ZmD11, increases seed size and quality in rice and maize. Plant Physiology and Biochemistry 160: 281–293.

Swigoňová Z, Lai J, Ma J, Ramakrishna W, Llaca V, Bennetzen JL, Messing J. 2004. Close Split of Sorghum and Maize Genome Progenitors. Genome Research 14: 1916– 1923.

Tong H, Chu C. 2012. Brassinosteroid Signaling and Application in Rice. Journal of Genetics and Genomics 39: 3–9.

Truernit E, Bauby H, Dubreucq B, Grandjean O, Runions J, Barthélémy J, Palauqui J-C. 2008. High-Resolution Whole-Mount Imaging of Three-Dimensional Tissue Organization and Gene Expression Enables the Study of Phloem Development and Structure in *Arabidopsis*. The Plant Cell 20: 1494–1503.

Vert G, Nemhauser JL, Geldner N, Hong F, Chory J. 2005. MOLECULAR MECHANISMS OF STEROID HORMONE SIGNALING IN PLANTS. Annual Review of Cell and Developmental Biology 21: 177–201.

Vilarrasa-Blasi J, González-García M-P, Frigola D, Fàbregas N, Alexiou KG, López-Bigas N, Rivas S, Jauneau A, Lohmann JU, Benfey PN, et al. 2014. Regulation of Plant Stem Cell Quiescence by a Brassinosteroid Signaling Module. Developmental Cell 30: 36–47.

Vriet C, Russinova E, Reuzeau C. 2012. Boosting Crop Yields with Plant Steroids. The Plant Cell 24: 842–857.

Wang Z-Y, Nakano T, Gendron J, He J, Chen M, Vafeados D, Yang Y, Fujioka S, Yoshida S, Asami T, et al. 2002. Nuclear-Localized BZR1 Mediates Brassinosteroid-Induced Growth and Feedback Suppression of Brassinosteroid Biosynthesis. Developmental Cell 2: 505–513.

Wang Y, Perez-Sancho J, Platre MP, Callebaut B, Smokvarska M, Ferrer K, Luo Y, Nolan TM, Sato T, Busch W, et al. 2023. Plasmodesmata mediate cell-to-cell transport of brassinosteroid hormones. Nature Chemical Biology 19: 1331–1341.

Wang Z-Y, Seto H, Fujioka S, Yoshida S, Chory J. 2001. BRI1 is a critical component of a plasma-membrane receptor for plant steroids. Nature 410: 380–383.

Warner CA, Biedrzycki ML, Jacobs SS, Wisser RJ, Caplan JL, Sherrier DJ. 2014. An Optical Clearing Technique for Plant Tissues Allowing Deep Imaging and Compatible with Fluorescence Microscopy. Plant Physiology 166: 1684–1687.

Wolf S, Mravec J, Greiner S, Mouille G, Höfte H. 2012. Plant Cell Wall Homeostasis Is Mediated by Brassinosteroid Feedback Signaling. Current Biology 22: 1732–1737.

Yamaguchi M, Fujimoto H, Hirano K, Araki-Nakamura S, Ohmae-Shinohara K, Fujii A, Tsunashima M, Song XJ, Ito Y, Nagae R, et al. 2016. Sorghum Dw1, an agronomically important gene for lodging resistance, encodes a novel protein involved in cell proliferation. Scientific Reports 6: 28366.

Yamamuro C, Ihara Y, Wu X, Noguchi T, Fujioka S, Takatsuto S, Ashikari M, Kitano H, Matsuoka M. 2000. Loss of Function of a Rice *brassinosteroid insensitive1* Homolog Prevents Internode Elongation and Bending of the Lamina Joint. The Plant Cell 12: 1591– 1605.

Yin Y, Wang Z-Y, Mora-Garcia S, Li J, Yoshida S, Asami T, Chory J. 2002. BES1 Accumulates in the Nucleus in Response to Brassinosteroids to Regulate Gene Expression and Promote Stem Elongation. Cell 109: 181–191.

Zhang C, Bai M, Chong K. 2014. Brassinosteroid-mediated regulation of agronomic traits in rice. Plant Cell Reports 33: 683–696.

