## Supplementary Figures for "Molecular and physiological characterization of brassinosteroid receptor BRI1 mutants in *Sorghum bicolor*"

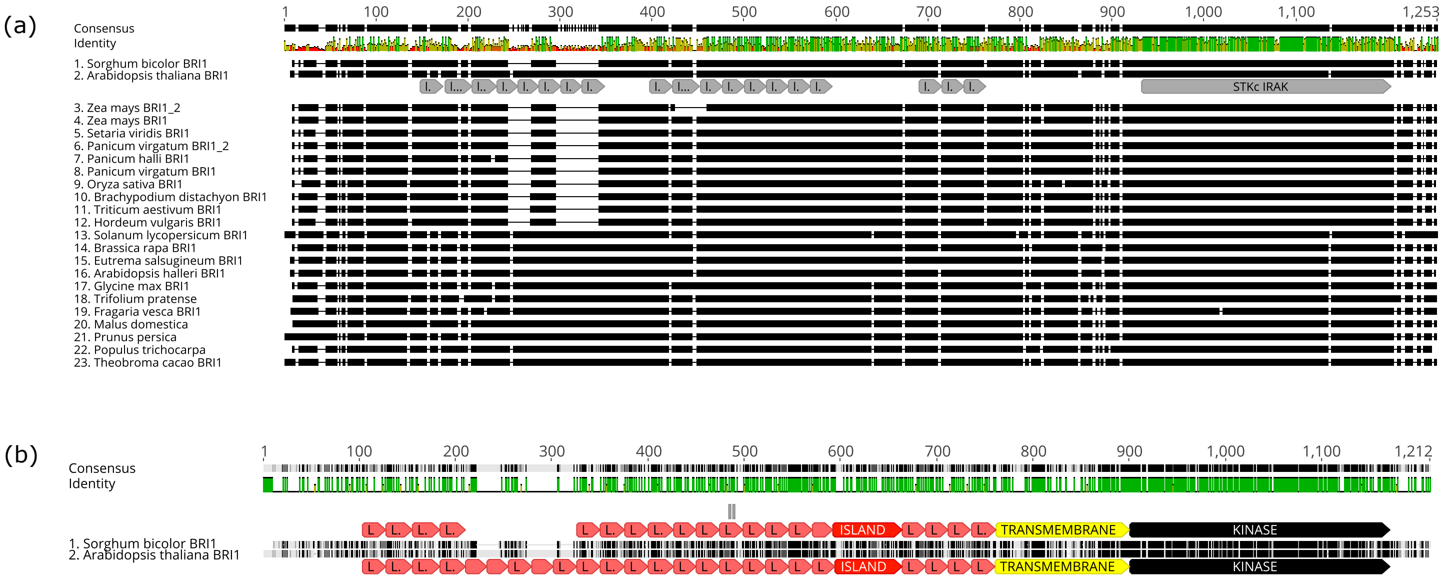


**Supplementary Figure 1:**
**A)** Sorghum and Arabidopsis BRI1 share a common sequence of kinase, transmembrane and island domain, suggesting functional conservation. Extracelular domain is mostly conserved with the exception of five LRR domains.
**B)** Alignment of multiple BRI1 sequences from different species showing a general conservation of the intracellular and extracellular domains.

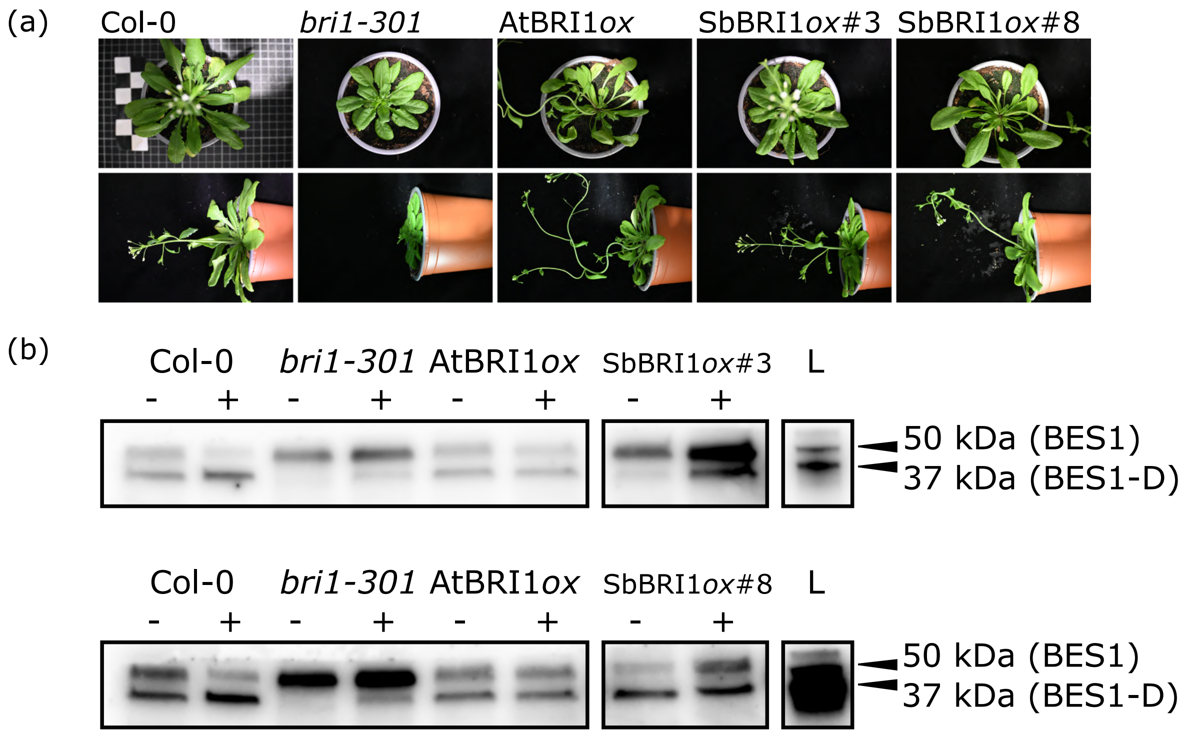


**Supplementary Figure 2:**
**(a)** SbBRI1 complements rosette size and flowering time in *bri1-301* Arabidopsis.
**(b)** BES1 dephosphorilation after BL treatment in SbBRI1 complemented lines of Arabidopsis *bri1-301* shows a functional conservation of BRI1 in sorghum.

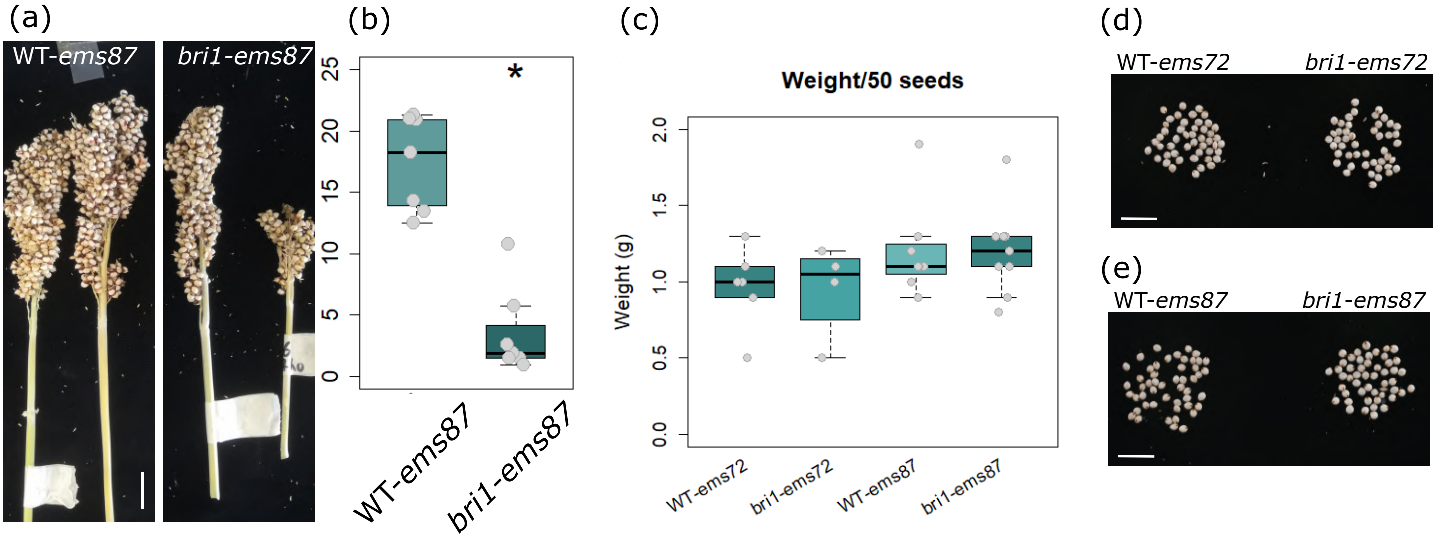


**Supplementary Figure 3:**
**(a, b)** *bri1-ems87* panicles present reduced yield when compared with their WT counterpart.
**(c)** seed weight of 50 seeds of WT-*ems* and *bri1-ems* mutants.
**(d, e)** size comparison of seeds of WT-*ems* and *bri1-ems* mutants.

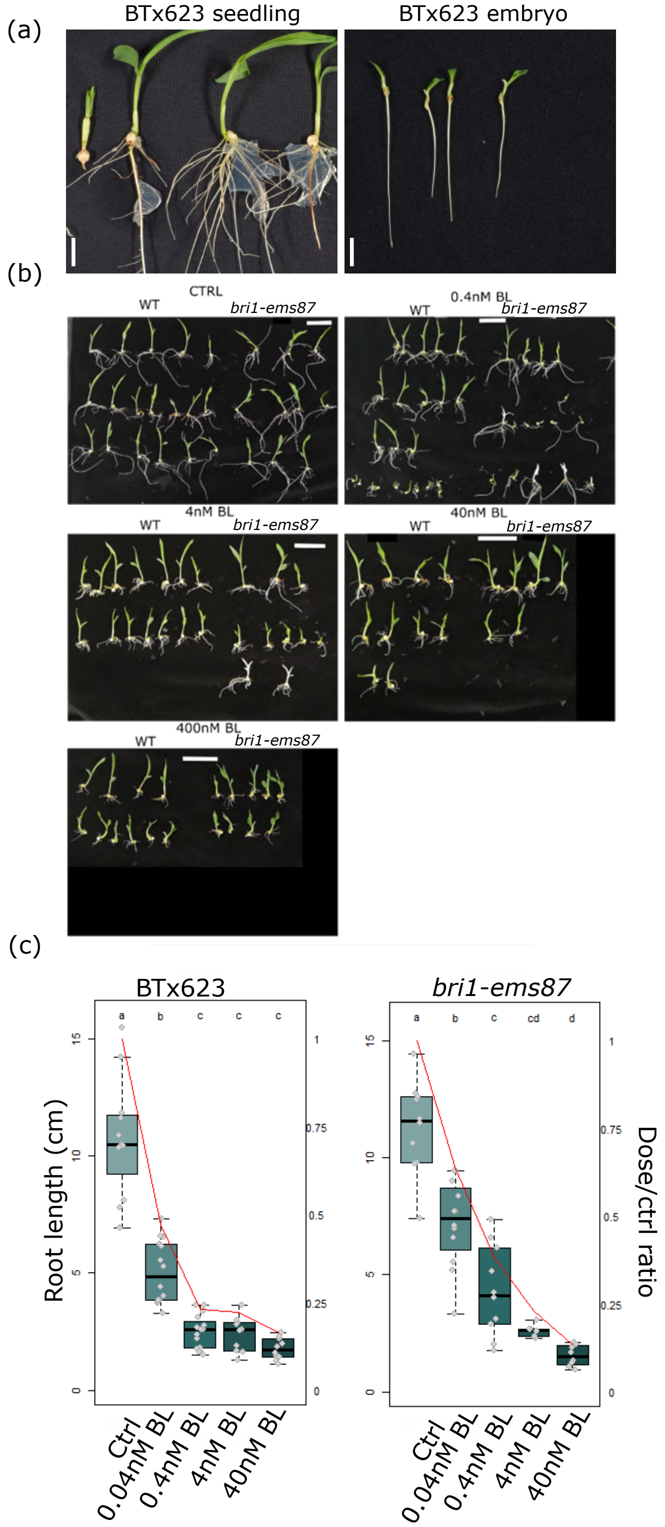


**Supplementary Figure 4:**
**(a)** Comparison of in vitro grown BTx623 from whole seedling vs from explanted embryo.
**(b)** Hydroponic grown sorghum seedlings show a insensitivity to BL of *bri1-ems87*.
**(c)** *bri1-ems87* sorghum root shows insensitivity to low concentrations of BL in terms of root length.

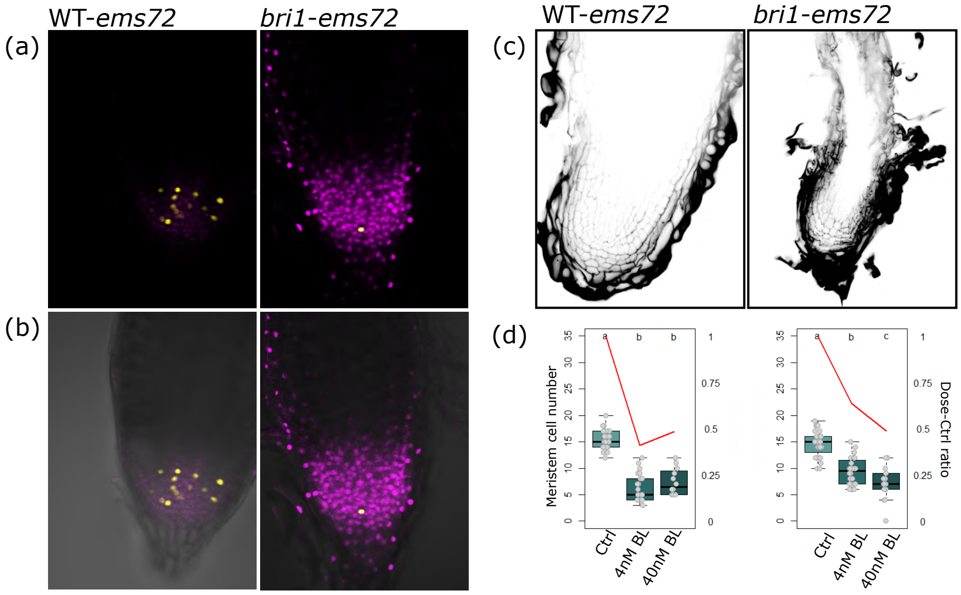


**Supplementary Figure 5:**
**(a)** Example at 40nm BL of root meristem exhaustion in WT-*ems72* and *bri1-ems72* (in yellow, dividing cells), **(b)** and with bright field bottom channel.
**(c)** Example of exhausted WT-*ems72* and *bri1-ems72* roots using mPS-PI show a great cell architecture disorganization in the root tip.
**(d)** At elevated BL concentrations, no differences were found between WT-*ems72* and *bri1-ems72*.

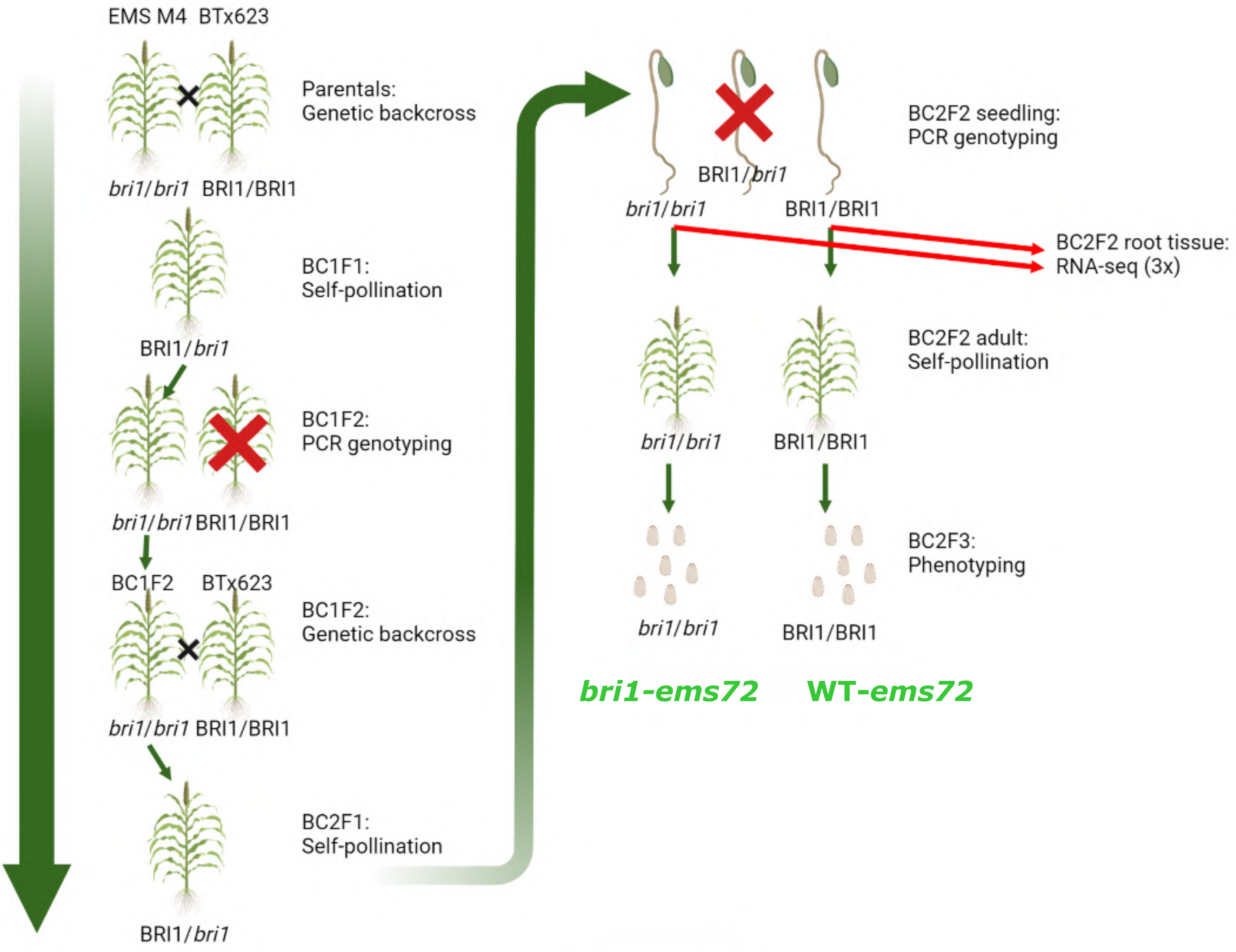


**Supplementary Figure 6: Diagram of backcrossing process to select our working seed lines.**
EMS-mutated M4 seeds of both *bri1* alleles from Jiao et al, 2016, were backcrossed against their unmutagenized parental BTx623 and PCR-genotyped for *bri1* mutations. Homozygous *bri1* line was selected for a second round of backcrossing and their progeny PCR-genotyped. F2 of this second backcross was PCR-genotyped using the aerial organs and used at seedling stage to perform RNA-seq of the root organ of both *bri1* and WT alleles of the BRI1 gene. Sibling progeny of both were grown and used to obtain F3 seeds for a comparative phenotypical analysis of BRI1.

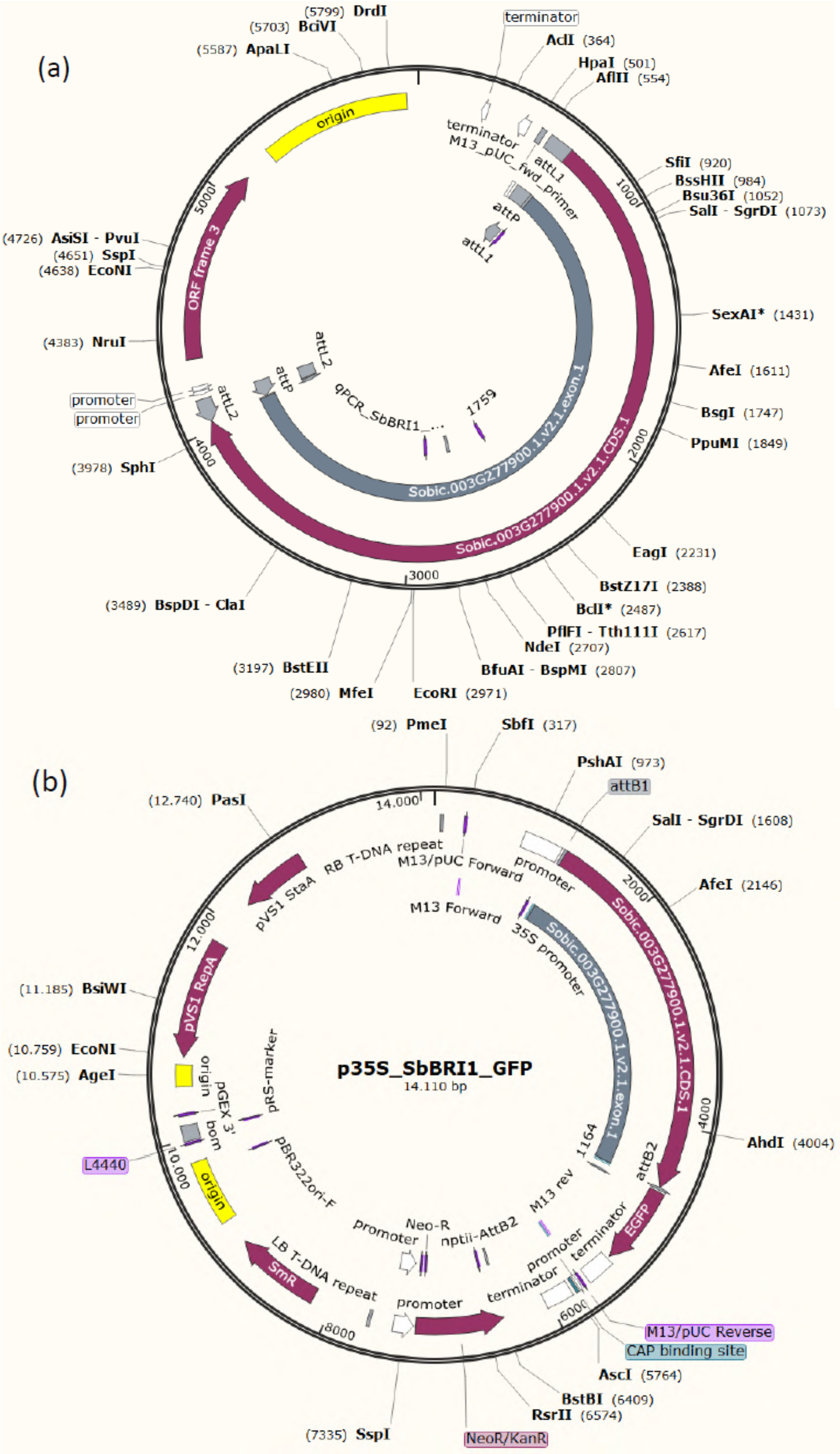


**Supplementary Figure 7:**
**(a)** After BP reaction, SbBRI1 coding sequence was cloned into pEN-L1L2 plasmid, giving pEN-L1L2-SbBRI1 entry clone.
**(b)** pEN-L1L2-SbBRI1 and the p4P1r-35S vector were LR-recombined with destination vector pGWB406 to obtain p35s:SbBRI1-GFP
